# The Lipid Droplet Knowledge Portal: A resource for systematic analyses of lipid droplet biology

**DOI:** 10.1101/2021.06.08.447431

**Authors:** Niklas Mejhert, Katlyn R. Gabriel, Natalie Krahmer, Leena Kuruvilla, Chandramohan Chitraju, Sebastian Boland, Dong-Keun Jang, Marcin von Grotthuss, Maria C. Costanzo, Jason Flannick, Noël P. Burtt, Robert V. Farese, Tobias C. Walther

**Affiliations:** Department of Molecular Metabolism, Harvard T.H. Chan School of Public Health, Boston, MA 02115, USA; Department of Cell Biology, Harvard Medical School, Boston, MA 02115, USA; Department of Medicine (H7), Karolinska Institutet, Huddinge, 141 86 Stockholm, Sweden; Howard Hughes Medical Institute, Boston, MA 02115, USA; Institute for Diabetes and Obesity, Helmholtz Zentrum München, 85764 Neuherberg, Germany; German Center for Diabetes Research, Neuherberg, Germany; Primary Pharmacology Group, Discovery Sciences, Pfizer Inc., Groton, CT 06340, USA; Program in Medical and Population Genetics, Broad Institute of Harvard and MIT, Cambridge, MA 02142, USA; Division of Genetics and Genomics, Boston Children’s Hospital, MA 02115, USA; Department of Pediatrics, Harvard Medical School, Boston, MA 02115, USA; Broad Institute of MIT and Harvard, Cambridge, MA, 02142, USA; Center on the Causes and Prevention of Cardiovascular Disease (CAP-CVD), Harvard T.H. Chan School of Public Health, Boston, MA 02115, USA

**Keywords:** inflammation, triacylglycerol, sterol ester, proteasome, C16orf54, MSRB3

## Abstract

Lipid droplets (LDs) are organelles of cellular lipid storage with fundamental roles in energy metabolism and cell membrane homeostasis. There has been an explosion of research into the biology of LDs, in part due to their relevance in diseases of lipid storage, such as atherosclerosis, obesity, type 2 diabetes mellitus, and hepatic steatosis. Consequently, there is an increasing need for a resource that combines large datasets from systematic analyses of LD biology. Here we integrate high-confidence, systematically generated data on studies of LDs in the framework of an online platform named the *Lipid Droplet Knowledge Portal*. This scalable and interactive portal includes comprehensive datasets, across a variety of cell types, for LD biology, including transcriptional profiles of induced lipid storage, organellar proteomics, genome-wide screen phenotypes, and ties to human genetics. This new resource is a powerful platform that can be utilized to uncover new determinants of lipid storage.

**HIGHLIGHTS:**

■ The LD-Portal is a resource combining datasets from systematic analyses in LD biology
■ The LD-Portal allows users to query genetic, proteomic, and phenotypic aspects of LD biology
■ The LD-Portal can be used to discover new facets of lipid storage and LD biology
■ A crucial function of *MSRB3* is uncovered in cholesterol ester storage in LDs

## INTRODUCTION

Lipid droplets (LDs) are phospholipid monolayer–bound organelles found in most eukaryotes and some prokaryotes. These organelles store neutral lipids, such as triacylglycerols (TGs) and cholesterol esters (CE), that can be used to generate metabolic energy or cell membranes. Specific proteins, including many important lipid metabolism enzymes (e.g., TG synthesis and degradation enzymes) bind to LD surfaces. Due to their important function in metabolism, alterations in LD biology are causal or implicated in diseases, such as lipodystrophy, atherosclerosis, obesity and related disorders (e.g., type 2 diabetes mellitus (T2D) and nonalcoholic fatty liver disease (NAFLD)). Moreover, alterations in LD metabolism are implicated in cancer, neurodegeneration, and immune function [1–5].

Despite the relevance of LDs to metabolic diseases, many aspects of their biology remain unclear, which has led to a recent surge of research into the biology of this organelle. In particular, systematic, unbiased approaches to studying LDs, including genome-wide screens to identify genes governing LD biology [6–9], and LD proteomics in different cells and tissues [6, 10–12], have been instrumental to progress in our understanding of LDs. However, the results from these various large-scale experiments are currently fragmented, limiting the integration and interrogation of data from various experiments. Specifically, unlike for other organelles, such as mitochondria [13], there is no comprehensive and scalable repository for integrating large data sets relevant to LD biology.

To address this deficiency and provide a resource for investigators of LD biology, we have created the *Lipid Droplet Knowledge Portal* (LD-Portal, lipiddroplet.org), an online resource that includes data from systematic research in LD biology. In this resource paper, we describe the initial version of the LD-Portal and, by highlighting several genes with phenotypes in these datasets that were previously not linked to LDs, we provide examples of how the LD-Portal can be used for discovering new facets of LD biology.

## RESULTS AND DISCUSSION

### Overview of the Lipid Droplet Knowledge Portal

A conceptual content map and detailed overview of the available data in the initial version of the LD-Portal are shown in **Figure 1**. The initial datasets integrated in the LD-Portal include a comprehensive dataset for RNA expression in cultured human macrophages (without and with lipid loading), LD proteomics for a variety of cell types, and a high-content imaging screen of genes governing LD biology in human cells [6]. In addition, the LD-Portal includes datasets for LD proteomics and phosphoproteomics of murine liver from mice fed chow or high-fat diets [10]. To enable efficient data mining of LD biology relevant to human physiology and disease, we integrated the LD-Portal data with human genetics data from the Common Metabolic Diseases Knowledge Portal (cmdkp.org). The LD-Portal resource allows both gene queries and phenotype-centric (“Gene Finder”) queries.

**Figure 1:**
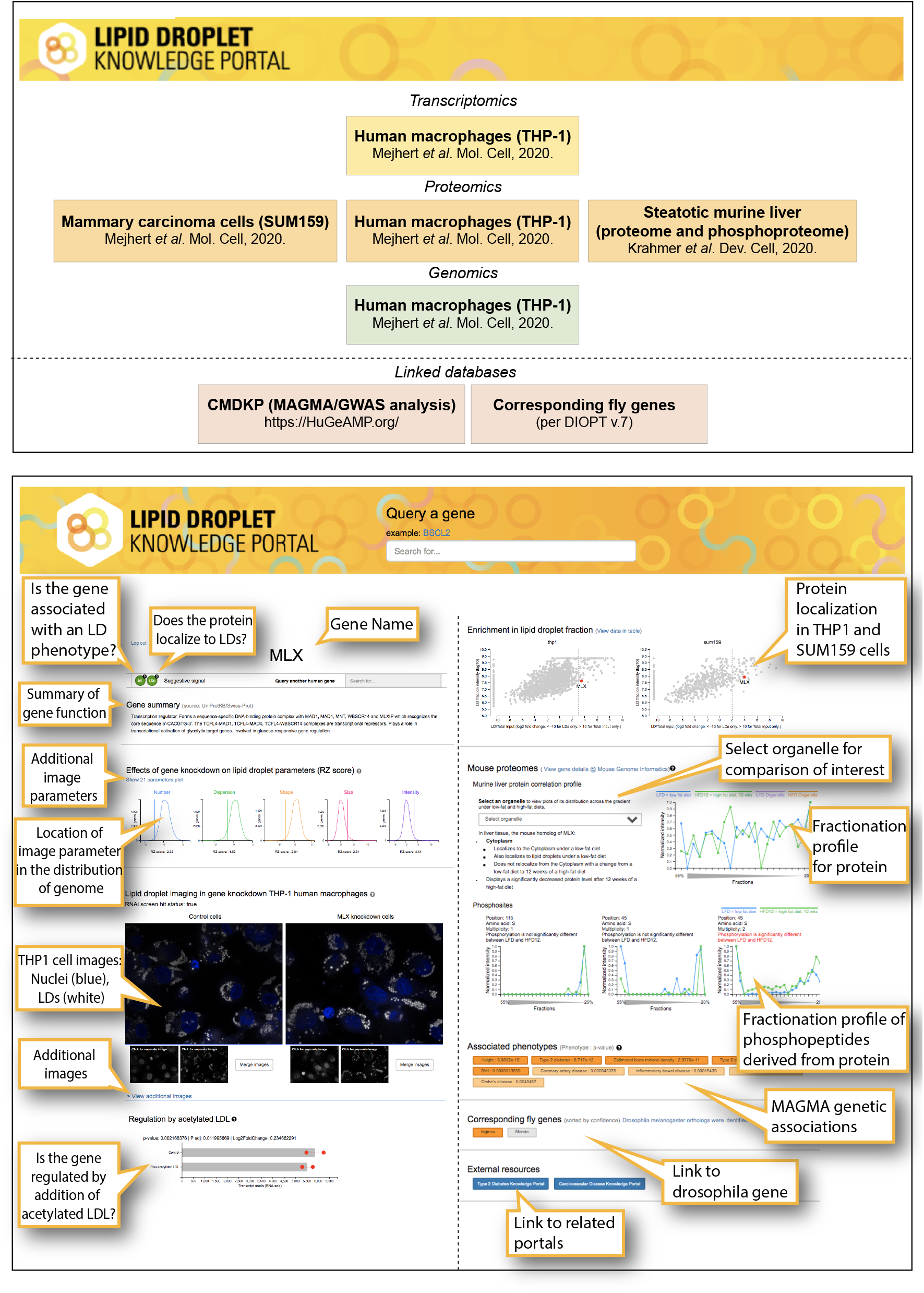
Overview of the Lipid Droplet Knowledge Portal. (A) Content of the LD-Portal. Original publications that contributed data to the initial version of the LD-Portal are listed. (B) Graphical summary of the LD-Portal interface with key to the data interface.

### Transcriptional Response to Increased Lipid Storage in Macrophages

Cells store excess lipids, such as fatty acids or sterols, as neutral lipids in LDs, a process that we named the “lipid storage response” (LSR, [6]). One component of the LSR is a rewiring of transcription to facilitate LD formation and lipid storage and utilization. To enable discovery of LSR mechanisms and integration with other aspects of LD biology, the LD-Portal includes information on gene expression changes in differentiated human THP-1 macrophages, under conditions promoting lipid storage, in this case by incubation of cells with lipoproteins that induce the formation of LDs. Cells were cultured with acetylated low-density lipoproteins (ac-Lipo) [6] that contain both CEs and TGs. Uptake and degradation of these lipoproteins in the endo-lysosomal pathway resulted in storage of both neutral lipids (**Figure 2A**). Induction of LDs containing both CE and TG enabled us to probe pathways for the intracellular storage of either neutral lipid; in contrast, incubation with oleic acid (OA) resulted in primarily TG accumulation in LDs (**Figure 2A**).

**Figure 2:**
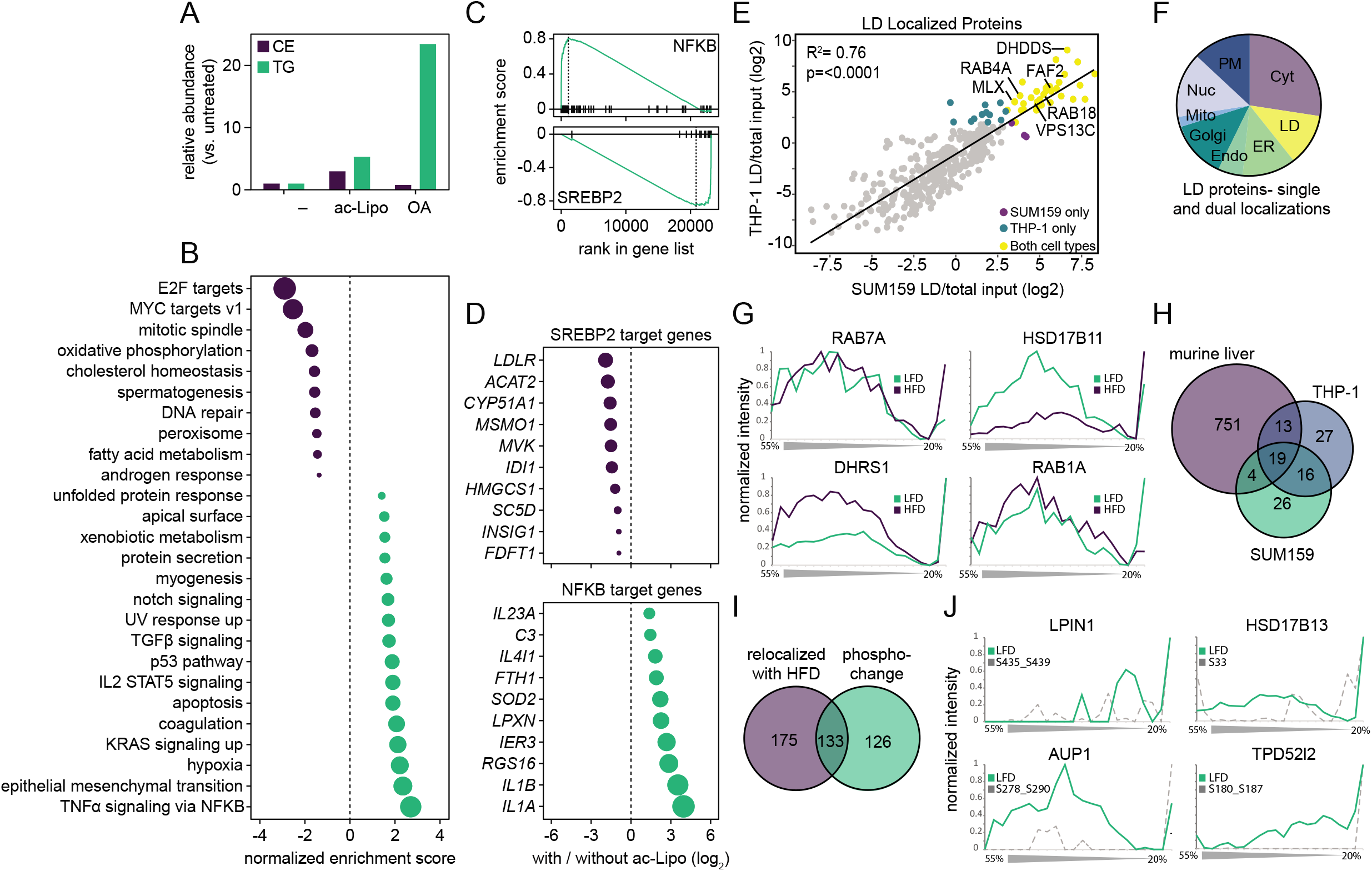
Data mining of the LD-Portal illustrating how lipid storage induction regulates metabolic and inflammatory pathways and protein localization to LDs. **(A)** Incubation with ac-Lipo induces TG and CE storage in THP-1 macrophages. Cellular lipid storage was induced by incubating differentiated THP-1 macrophages in the absence/presence of ac-Lipo (100 μg/mL) or OA (0.5 mM) for 1 day, followed by determination of lipid composition by thin layer chromatography. Results from one experiment are shown. **(B-D)** Metabolic and inflammatory pathways are regulated by macrophage lipid storage. Transcriptional profiles of THP-1 macrophages incubated in the presence/absence of ac-Lipo (50 μg/mL) for 2 days were determined using RNA sequencing. (B) Pathways regulated by lipid storage were identified using gene set enrichment analysis based on the hallmark gene set. (C) As a proxy for transcriptional activities, validated SREBP2 and NFκB target genes were ranked across the RNA sequencing results and enrichment scores calculated. (D) Top 10 regulated SREBP2/NFκB target genes from panel C are displayed. Results are based on two replicates per condition, and in panels (B) and (D), the size of each circle is scaled to match the absolute value of the respective x-axes. **(E)** Multiple proteins are localized to LDs in both SUM159 and THP-1 cells. The log_2_ foldchange of the LD fraction/total input was plotted for proteins common to the LD fraction of both SUM159 and THP-1 cells. Intensity cutoffs of 3.066 and 2.02 were used for THP-1 and SUM159 cells, respectively. Proteins that were over the threshold in both cell types are labeled in yellow, and proteins that fulfilled the criteria in only one of the cell types are highlighted in purple (SUM159) and blue (THP-1). **(F)** In the mouse liver PCP, 787 proteins localized to LDs: 94 had a unique LD localization (yellow) and others localized to LDs and the indicated second organelle. **(G)** PCP profiles of RAB7A, HSD17B11, DHRS1, and RAB1A show a strong signal in LD fractions, indicating a relocalization of the protein to LDs under HFD conditions. **(H)** Venn diagram showing overlap of proteins from murine liver, SUM159 and THP-1 LD proteomes. **(I)** Venn diagram of murine liver PCP LD proteins that relocalized with HFD and LD proteins with phosphorylation changes with HFD. **(J)** Protein profile of the LD proteins LPIN1, HSD17B13, AUP1, and TPD52l2 overlaid with the profile of an LD localization-specific phosphorylation. Abbreviations: ac-Lipo, acetylated apolipoprotein B-containing lipoprotein; CE, cholesterol ester; OA, oleic acid; TG, triacylglycerol.

Our RNA sequencing studies of the LSR in THP-1 cells showed pronounced changes in gene expression for 2414 genes (1289 up-regulated, and 1125 down-regulated, adj. p-value<0.01) after culturing cells in the presence of ac-Lipo [6]. New analyses of these data with gene set enrichment showed expected changes, such as the downregulation of cholesterol homeostasis genes and the induction of expression of the unfolded protein response and inflammatory genes (**Figure 2B**). However, genes in a large number of other categories were also significantly altered during the LSR of this cell type. For many of these genes, the relationship of changes in their expression to ac-Lipo treatment is currently unknown. Further analyses of this dataset revealed that SREBP2 target genes were predominantly downregulated, explaining the changes in cholesterol homeostasis gene expression, and NF-kB target genes were upregulated, explaining increased expression of inflammation-related genes (**Figure 2C-D**). This dataset, accessible on the LD-portal interface, therefore provides a rich resource for probing the cellular response to lipid loading in macrophage cells.

### Lipid Droplet Proteomes of Human Cells and Murine Liver

The LD-Portal also includes data on the subcellular localization of proteins and particularly highlights the potential of proteins to localize to LDs under different conditions. To collect comprehensive information on the LD proteome of several model systems, we integrated data from the proteomic analyses of LDs from human THP-1 macrophages, human SUM159 triple-negative breast cancer cells (used frequently to study LD biology) (both datasets originally published in [6]), and large-scale *in vivo* murine proteomic and phosphoproteomic organellar-localization atlas (dataset originally published in [10]). These data are now collectively available for analysis on the LD-Portal.

For the SUM159 and THP-1 cultured cell lines, we measured the enrichment of proteins in the LD fraction compared to the total input fraction to identify proteins enriched on LDs. Based on the enrichment of known LD proteins identified in Bersuker *et al*. [14], we calculated confidence intervals for the enrichment score and used these as cut-offs for assigning a protein to LD localization. For THP-1 cells, a total of 5801 proteins were detected in the whole-cell lysate, with 1412 proteins in the LD fraction and 75 enriched there (fold-change threshold, 3.07). For SUM159 cells, 5708 proteins were detected in whole-cell lysate, with 629 proteins in the LD fraction, and 64 enriched in this fraction (fold-change threshold, 2.02). To assess the robustness of the THP-1 and SUM159 proteomes, we compared the protein intensities in the LD fraction of THP-1 macrophages to that of SUM159 cells and found that these datasets were well correlated (R^2^=0.76, p<0.0001), with 35 specific LD proteins in common, including MLX, VPS13C, RAB18, FAF2, RAB4A, and DHDDS (**Figure 2E**). A list of the LD-enriched proteins found in both cell types is provided in **Table S1**. Of the 35 LD proteins common to both cells, 30 were also reported in an APEX2-mediated screen to identify LD proteins in U2OS or HuH7 cells [12, 14]. However, we identified 42 additional LD proteins, including PLIN4, CHP1 and LPCAT2.

The LD-Portal also includes data on subcellular protein localization based on protein correlation profiling for the majority of proteins across organelles in C57BL/6J murine liver [10]. These studies were performed in mice fed chow or high-fat diets (HFD), and by examining proteins across different cell fractions, they revealed how nutrient overload leads to organellar reorganization [10]. Of the 6163 proteins quantified across cellular fractions, 5878 gave reproducible profiles for organelle assignment. We found diet-dependent re-localization for 901 proteins, and protein expression changes for 258. We assessed the reproducibility of this dataset by calculating Pearson correlation of profiles, derived from the same biological conditions and between different diets, with Pearson correlations of 0.86 and 0.78 for protein levels and relocalization patterns, respectively (**Figure S1A**). For the murine liver samples, 787 protein profiles showed a characteristic peak in the top fraction after organelle separation by density centrifugation, indicating localization on the LDs or in LD-associated membranes. Most of these proteins localized to multiple organelles, and only 94 had a unique LD localization (**Figure 2F**). Of the 787 LD proteins, 308 showed a significant profile shift under HFD feeding. For instance, the proteins RAB7A, HSD17B11, DHRS1 and RAB1A undergo HFD-induced relocalization to LDs (**Figure 2G**).

We also utilized the LD-portal data to compare the LD proteins detected in THP-1 and SUM159 cells and murine liver. Based on these data, we identified 19 LD proteins that were common to all three datasets (**Table S1**). These include well-known LD proteins (e.g., AUP1, ACSL3, MLX, and PCYT1A) but also several for which data on LD localization was previously sparse (e.g., VPS13C (a lipid trafficking protein) and LSS (lanosterol synthase)). An additional 33 proteins were found to be in common to at least two datasets (**Figure 2H**), including well-known LD proteins (e.g., PLIN4) and proteins previously not established as LD proteins (e.g., NRZ tethering protein NBAS and TACC1 (**Table S1**). Notably, the function in this organelle is unclear for many of the proteins that were reproducibly and robustly identified in LD fractions. We anticipate these proteomic datasets will open numerous new lines of investigation.

### LD Proteins of Murine Liver That Are Phosphorylated

The LD-Portal also includes comprehensive data on the localization of phosphorylated forms of proteins within murine liver [10]. From 24,524 phosphosites, 11,712 gave reproducible profiles, and 1676 phosphorylation levels changed with HFD after normalization to protein levels. Analyzing specifically the LD proteins, 3037 had partial and 229 had unique LD localization, as assigned by support vector machine-based organelle assignments. Among all proteins targeted from other compartments to LDs under HFD, almost half of the re-localizations (133) were accompanied with phosphorylation changes (**Figure 2I**), indicating that this might be an important regulatory mechanism for targeting of specific proteins to LDs.

Overlaying the protein and phosphosite profiles enabled identification of localization-specific phosphosites and phosphorylation events that are independent of protein localization. For example, the profile for AUP1, a protein identified as an LD protein [15] shows dual LD and ER localization (**Figure 2J**). Yet, the same protein had phosphosites (S287/S290) that appeared only in the LD part of the profile, indicating that this site is phosphorylated only in the LD fraction and not the ER pool of the protein. Similarly, S33 of HSD17B13, S180_S187 of TPD52l2, and S180_S187 of LPIN1 had LD-specific phosphosignatures (**Figure 2 J**). These examples illustrate how the portal can be used in future studies to identify phosphosites that might regulate protein localization to LDs.

### Genome-Perturbation Screen for Lipid Droplet Phenotypes in Human Cells

The LD-Portal additionally features a large dataset from a high-content, imaging-based genome perturbation screen [6]. In this screen, LDs were induced by ac-Lipo and stained with BODIPY, and LD information was collected using automated imaging and extraction of multiple image parameters at the single-cell level by image segmentation [6] (**Figure 3A**). With this pipeline, we disrupted expression of essentially all genes one-by-one in triplicate experiments and analyzed the effects on LDs in THP-1 macrophages. This dataset was utilized, for example, to discover that the MLX family of transcription factors (e.g., MLX, MLXIP, MLXIPL/ChREBP) bind LDs and modulate their transcriptional activity [6].

**Figure 3:**
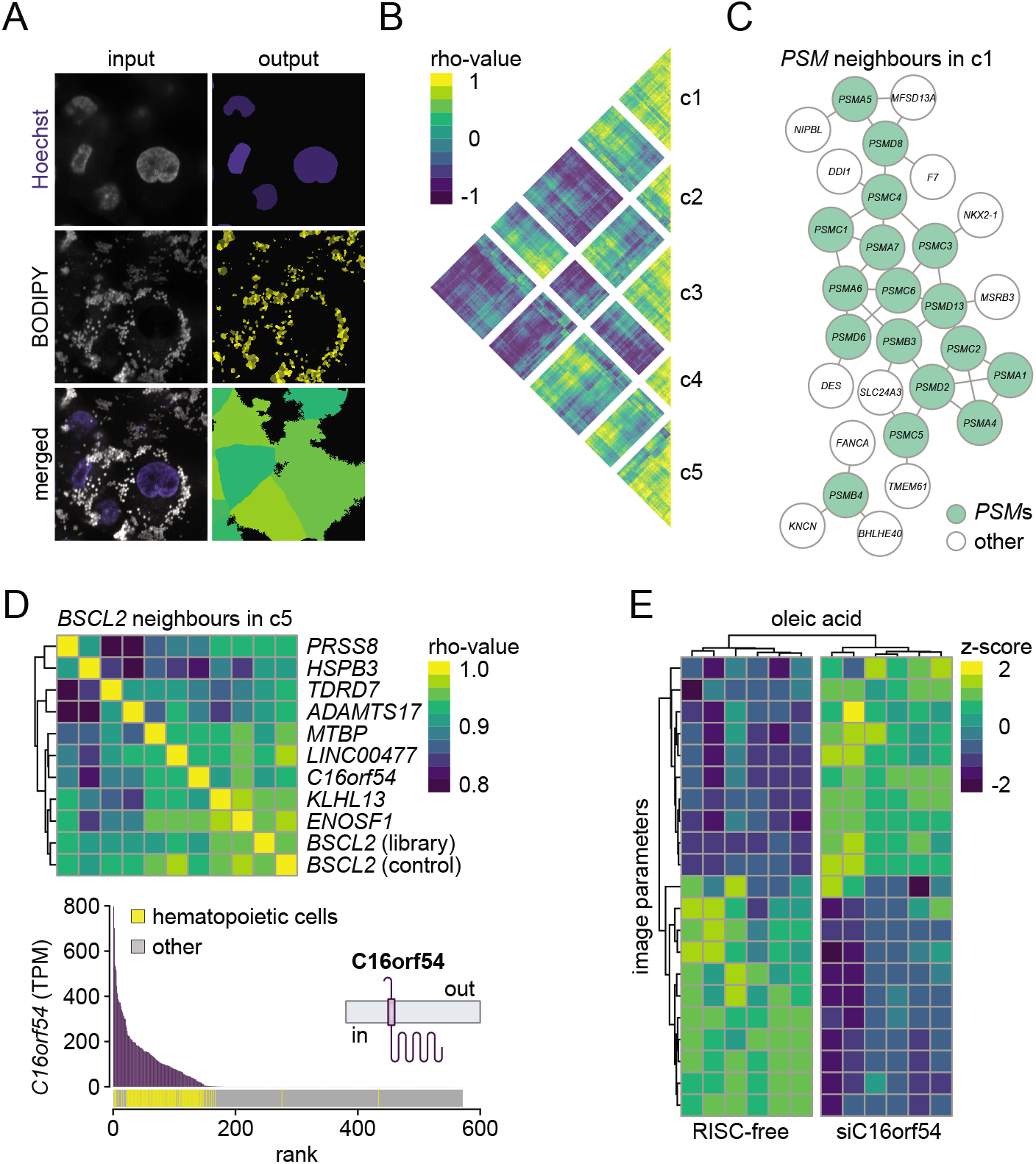
Clustering of RNAi screen image results identifies classes of hits with similar LD morphologies. **(A)** Image analysis extracts LD information at the single-cell level. After lipid storage induction by ac-Lipo, nuclei and LDs of THP-1 macrophages were stained using Hoechst and BODIPY, respectively. Images were acquired using a high-throughput confocal microscope, and image analyses were performed using CellProfiler. Segmented nuclei, LDs and cells are shown in the output column. **(B)** Five classes of macrophage lipid determinants. RNAi screen hits (n=558) were pair-wise correlated, based on image information, and the resulting matrix was classified (c1-5) by hierarchical clustering. c1 contains predominantly proteasome genes. c5 is the *BSCL2*/seipin cluster. **(C)** Knockdown of proteasomal subunits results in small and dispersed LDs. Network displaying the 17 proteasomal subunits and their closest neighbors identified in cluster 1. The three closest neighbors of each proteasomal gene in cluster 1 were extracted from the correlation matrix presented in panel (B) and added to the network. Genes are presented as nodes, and top three neighbors are connected by edges. **(D-E)** Depleting *C16orf54* and *BSCL2* results in similar macrophage LD morphology. (D) In the upper panel, correlation scores for the 10 closest neighbors of *BSCL2* in cluster 5 are displayed. *BSCL2* occurs two times: once from the genome-wide library (library) and once as a median score of all BSCL2 control wells present on each plate (control). In the lower panel to the left, *C16orf54* gene expression data across 571 human cells and tissues were extracted from the FANTOM5 database. *C16orf54* transcript abundance was ranked from high to low, and samples from hematopoietic cells were highlighted. In the lower panel to the right, transmembrane helices in C16orf54 were predicted using TMHMM Server v. 2.0. (E) After incubations with oleic acid (0.5 mM) for 1 day, LD image analysis of RISC-free control and siC16orf54 transfected THP-1 macrophages was performed. Each feature was scaled, and the resulting matrix was plotted as a heatmap. Results are based on six replicates. Abbreviations: c1-5, cluster 1-5; LD, lipid droplet; PSMs, proteasomal subunits; TPM, tags per million.

Our analyses of the data from this screen yielded 21 non-redundant image parameters that describe LD size, number, dispersion, shape, and intensity in the screen images ([6] and **Figure 1, bottom panel**). Using tools from the LD-Portal, we performed additional analyses of these screen data. When we clustered genes with similar effects on LD parameters, we identified clusters containing genes with similar biological functions (**Figure 3B, Table S2**). For example, cluster 1 (c1) contains 17 genes involved with proteasome function (**Table S2**). The similarity of phenotype associated with different proteasomal (PSM) subunit genes is also apparent in a network plot that shows the genes with the most similar phenotype for each proteasome subunit in cluster 1 (**Figure 3C**). By this analysis, each of the *PSM* genes, except *PSMB4*, are interconnected. Finding LD phenotypes for disruptions of PSM genes is consistent with our previous RNAi screen in *Drosophila melanogaster* S2 cells [7]. In addition, 12 more genes are part of this proteasome network. Among these, several genes (e.g., *DDI1*) have been directly implicated in the ubiquitin-proteasome system [16]. In addition, *TMEM61*, which encodes an unknown ER protein, was reported in human genetic datasets as being highly associated with cholesterol metabolism and has limited homology to scavenger receptors (HHPRED, [17]).

For the four other clusters (c2, c3, c4, and c5) highlighted in **Figure 3B**, the biological underpinnings for similar LD phenotypes are unknown and not readily apparent. Nonetheless, these clusters are likely to be informative for LD biology. For instance, independent replicates of *BSCL2*, encoding the protein seipin—a key factor in LD formation [18–22]—clustered tightly, validating the approach, and this phenotype (found in cluster 5) identified several genes whose depletion phenotypes were highly similar (**Figure 3D**). The phenotype for *BSCL2*/seipin depletion was also similar to that of *LDAF1* depletion (not classified as a “hit” by stringent criteria, but shown in **Figure S2**), and the proteins encoded by these two genes function together in an LD formation complex [18]. Another hit tightly correlated with *BSCL2*/seipin is the uncharacterized *C16orf54* (**Figure 3D, upper panel**). This open-reading frame is predicted to encode a protein with one transmembrane domain and is expressed highly in hematopoietic cells (**Figure 3D, lower panels**). Thus, the encoded protein may be functionally related to BSCL2/seipin, possibly with a functional role in blood cells. As seipin regulates LD formation induced by fatty acid supplementation, we tested if knockdown of *C16orf54* in THP-1 macrophages changed LD morphology when incubating the cells with oleic acid. Our results show that *C16orf54* is required for normal lipid storage induced by lipoproteins and fatty acids (**Figure 3E**). Furthermore, our initial analyses of these screen data reveal groups of genes with similar LD depletion phenotypes and suggest that further mining of correlated genes may yield many mechanistic discoveries of machinery or pathways affecting LD biology.

### Data Mining of the LD-Portal Identifies MSRB3 as a Determinant of Cholesterol Ester Storage

As an example of how the LD-Portal can be mined for new insights to LD biology, we performed a secondary screen of randomly selected genes in which we compared LD phenotypes in response to gene knockdowns when LD formation was driven by cholesterol (via culture with ac-LDL) and fatty acids (via culture with oleic acid) (**Figure 4A**). This screen revealed that some genes, such as *BCSL2/Seipin*, exhibited robust phenotypes for either culture condition. Other genes, such as C9orf16, were more important for storage of one of the excess lipids.

**Figure 4:**
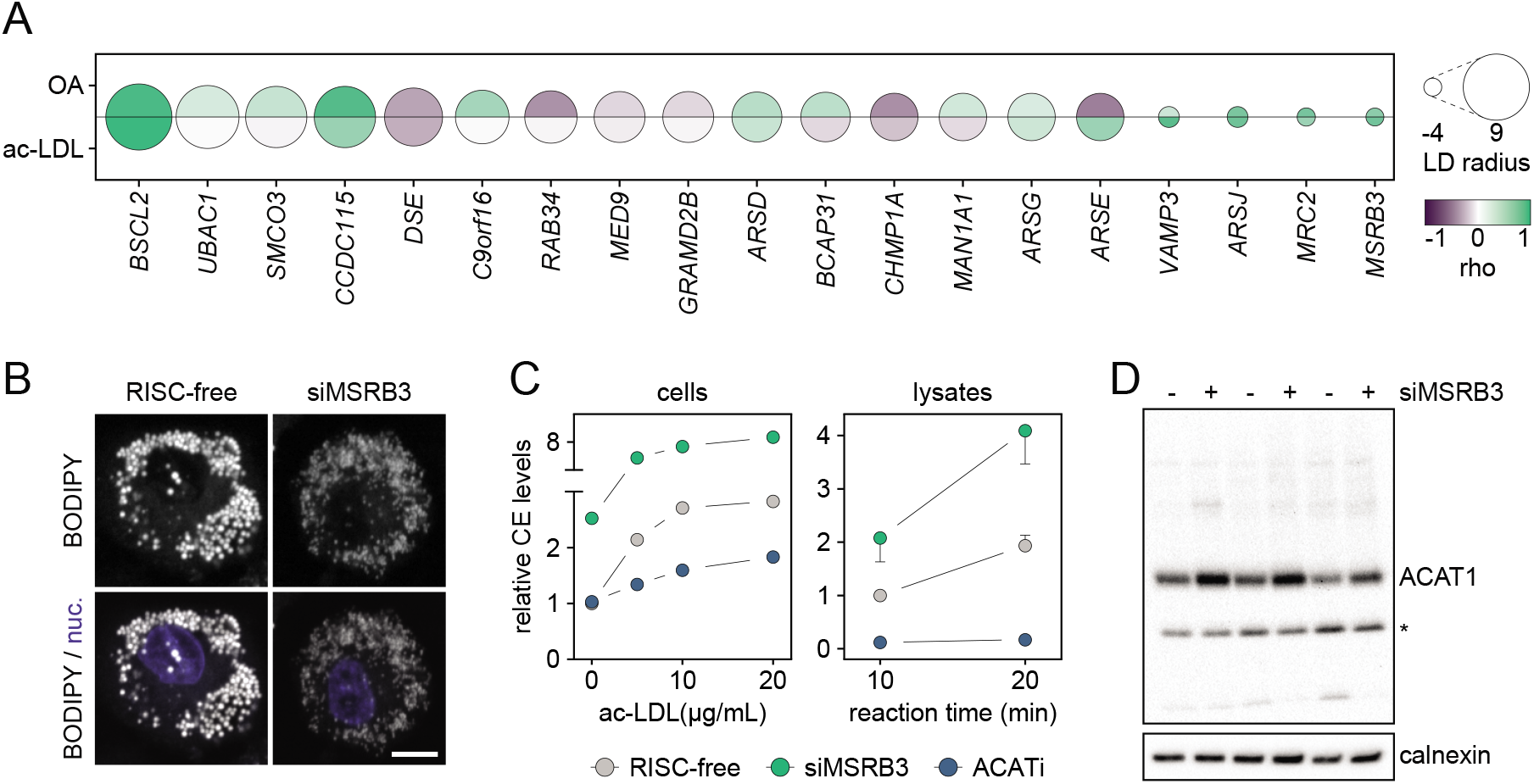
Secondary screening identifies genetic determinants of cholesterol ester versus triacylglycerol storage and identifies MSRB3 as a regulator of cholesterol storage. **(A)** Randomly selected genes were re-screened in THP-1 macrophages with ac-LDL or OA. The size and color of the circle are proportional to the effect on LD size in the original screen and reproducibility in the two secondary screens, respectively. **(B)** Macrophage *MSRB3* knockdown results in small and dispersed LDs. Representative confocal images of RISC-free or siMSRB3 transfected macrophages from the original RNAi screen. Scale bar, 5 μm. **(C)** Cholesterol esterification is increased in *MSRB3*-depleted macrophages. THP-1 macrophage cholesterol esterification assays were performed in live cells (left panel) or lysates (right panel) 3 days post-transfection with siRNAs targeting RISC-free or siMSRB3. ACAT inhibition was used as a control for the assays, and levels of extracted radiolabeled CE were determined by thin layer chromatography. In the left panel, endogenous production and exogenous uptake of cholesterol were reduced in cells by culturing them with compactin and without FBS, respectively. Subsequently, ac-LDL was added to the media for 7 hours out of which the 2 last hours were in the presence of radiolabeled OA. One representative experiment is shown. In the right panel, radiolabeled cholesterol was added to cell lysates, and CE formation was allowed for the indicated reaction times. Results are based on four replicates. **(D)** ACAT1 protein levels are increased in *MSRB3*-depleted macrophages. Protein levels of ACAT1 and calnexin were determined by western blotting in THP-1 macrophages 3 days posttransfection of RISC-free or siMSRB3 siRNAs. In addition to the ACAT1 band (approximately 45 kDa), an unspecific band marked by an asterisk was detected (approximately 30 kDa). Results display three independent experiments. Abbreviations: ACATi, ACAT inhibition; ac-LDL, acetylated low-density lipoprotein; nuc, nuclei; OA, oleic acid.

This secondary screen identified *MSRB3* as a gene with a striking depletion phenotype characterized by small and dispersed LDs (**Figure 4B**). In the LD-Portal (see below), *MSRB3* was associated with changes in type 2 diabetes (T2D) (p=3.8e-5) and adiponectin levels (p=7.7e-5), among other traits. *MSRB3* encodes an ER methionine sulfoxide reductase of unclear function. Mutations in human *MSRB3* lead to deafness, and it has been associated with progression of renal clear cell carcinoma, gastric cancer and Alzheimer’s disease [23–27] Most often, sulfoxide reductases are thought to help to maintain protein folding, structure and activity. To determine the biochemical basis of the LD phenotype, we investigated synthesis of cholesterol esters in cells and lysates depleted for *MSRB3* (**Figure 4C**). Cholesterol ester synthesis was significantly increased when *MSRB3* was absent. Increased levels of the main cholesterol ester synthesis enzyme in THP-1 macrophages, *SOAT1*/ACAT1, were also seen (**Figure 4D**). These findings suggest MSRB3 is required to control ACAT1 transcription, protein turnover, or activity, and this hypothesis can now be investigated in mechanistic detail to determine how a methionine sulfoxide reductase affects cellular cholesterol metabolism.

### Integration of Lipid Droplet Biology Datasets with Human Genetics

The LD-Portal also contains human genetic gene and gene-set association analyses for many complex traits, calculated using the *Multi-Marker Analysis of GenoMic Annotation* (MAGMA) algorithm [28]. To explore these connections, we determined if our datasets of LD proteins or gene hits associated with an LD-phenotype had preferential association scores in the MAGMA dataset. We defined a subset of MAGMA traits as metabolically associated, including BMI, coronary artery disease, child obesity, inflammatory bowel disease, T2D, waist circumference, visceral adipose tissue volume, and total cholesterol, ALT, adiponectin, oleic acid, palmitic acid, palmitoleic acid, fasting glucose, fasting insulin, HDL-cholesterol, LDL-cholesterol, and TGs.

We examined MAGMA metabolic association scores for 558 RNAi THP-1 screen hits, 3220 genes whose RNA expression levels were modulated by ac-Lipo, and 755 LD-localized proteins, and we found a number of expected associations in each set of genes (**Figure 5A**). For example, we detected strong associations of *APOB* and *APOE* with cholesterol and LDL phenotypes; scavenger receptor class B type 1 (*SCARB1*) with HDL cholesterol; insulin growth factor 1 (*IGF1*) with fasting insulin, and *FTO* (associated with fat mass and obesity) with BMI [29–31]. Within each of these datasets, we detected significantly more associations with metabolic traits than expected for a random sample of genes (**Figure 5B**) (p-values=0.025, 6.72e-5, and 3.09e-5, respectively).

**Figure 5:**
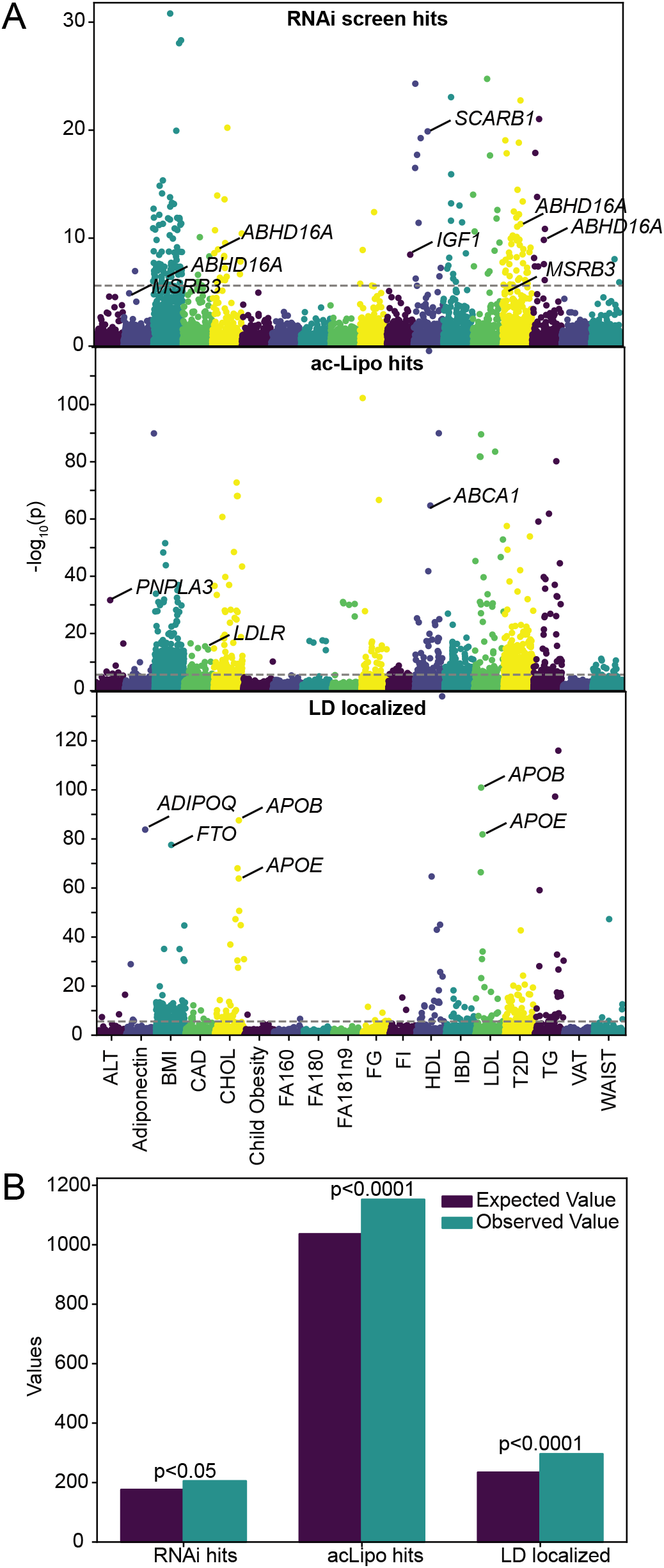
The LD portal screen gene sets are associated with human genetic traits. **(A)** Metabolic associations (-log_10_ p values) of genes that were either RNAi screen hits, ac-Lipo screen hits, or LD localized proteins (for each SUM159, THP-1 and murine proteomes) were analyzed for 19 metabolic phenotypes. A significance threshold value of p = 2.5e-6 was used (dotted line). **(B)** Chi-squared results of RNAi screen hits, ac-Lipo screen hits, and LD localized proteins that have significant associations with metabolic phenotypes vs random selection. Chi-squared statistics= 5.00, 15.89, and 17.36, respectively. Abbreviations: ALT, alanine transaminase; BMI, body mass index; CAD, coronary artery disease; CHOL, cholesterol; ChildObesity, child obesity; FA160, palmitic acid; FA180, palmitoleic acid; FA181n9, oleic acid; FG, fasting glucose; FI, fasting insulin; HDL, HDL cholesterol; IBD, inflammatory bowel disease; LDL, LDL cholesterol; T2D, type 2 diabetes; TG, triglyceride; VAT, visceral adipose tissue volume; WAIST, waist circumference

Analyzing the hits of the genome-perturbation screen in THP-1 cells, we detected a number of highly significant associations with human metabolic traits. For instance, we identified *ABHD16A* associated with five different metabolic traits, including BMI, cholesterol levels, T2D, TGs, and ulcerative colitis. We plotted the MAGMA score percentile for each gene, highlighting *ABHD16A* for nine total traits (**Figure S3**). Eight traits had significant associations, indicating a p-value less than 2.5 E-6 (6.4 on a log_10_ scale). The function of ABHD16A is not well-understood, but molecularly it encodes a phosphatidylserine hydrolase of the ER [32]. Our data suggest that modulating phosphatidylserine levels is important to maintain normal LDs, and interference with normal phosphatidylserine levels can lead to metabolic complications.

### Future Outlook

The LD-Portal provides a rich open-source platform for mining biological databases related to LD biology. The current version of the LD-Portal provides several searchable databases that can be mined to query genes or phenotypes and discover new connections for further mechanistic exploration. Additionally, integration of LD-Portal data with other platforms, such as human genetic MAGMA data from the Common Metabolic Diseases Knowledge Portal, allows filtering of queries to discern connections with human disease. In this description of the LD-Portal resource, we highlighted several examples, based on our initial analysis of the data sets, illustrating how mining of the LD-Portal resources will undoubtedly advance discoveries in LD biology.

The LD-Portal is scalable, allowing for integration of data from various sources, including genome perturbation screens, proteomic studies, gene expression analyses, and in the future, lipidomics or metabolomics data. The database is designed to allow integration of datasets from the community (with contact information and links on the LD-Portal website), which will enable an expanding platform of data relevant to LD biology. With many important links of lipid metabolism and LDs to prevalent public health problems, such as obesity, hepatic steatosis and NAFLD/NASH, and cardiovascular disease, the LD-Portal will also enable new insights into the basic biology of important genetic risk factors for these diseases.

## ACKNOWLEDGMENTS

We thank Drs. Ta-Yuan and Catherine Chung-Yao Chang (Department of Biochemistry, Dartmouth Medical School) for kindly providing a custom-made ACAT1 antibody. This work was supported in part by the Howard Hughes Medical Institute and NIH grant R01DK124913 (to R.V.F. and T.C.W.).

## AUTHOR CONTRIBUTIONS

N.M., K.R.G., N.K., R.V.F., and T.C.W. started the project and wrote the first version of the manuscript. N.M., K.R.G., N.K., and L.K. generated the screen and proteomics data based on SUM159 cells, THP-1 macrophages and mouse liver. C.C. and S.B. performed the cholesterol esterification assays in lysates and cells, respectively. The portal was built by D-K.J., M.v.G., M.C.C., J.F., and N.P.B. All co-authors read, commented, and approved the final version of the manuscript.

## SUPPLEMENTARY INFORMATION

**Figure S1:**
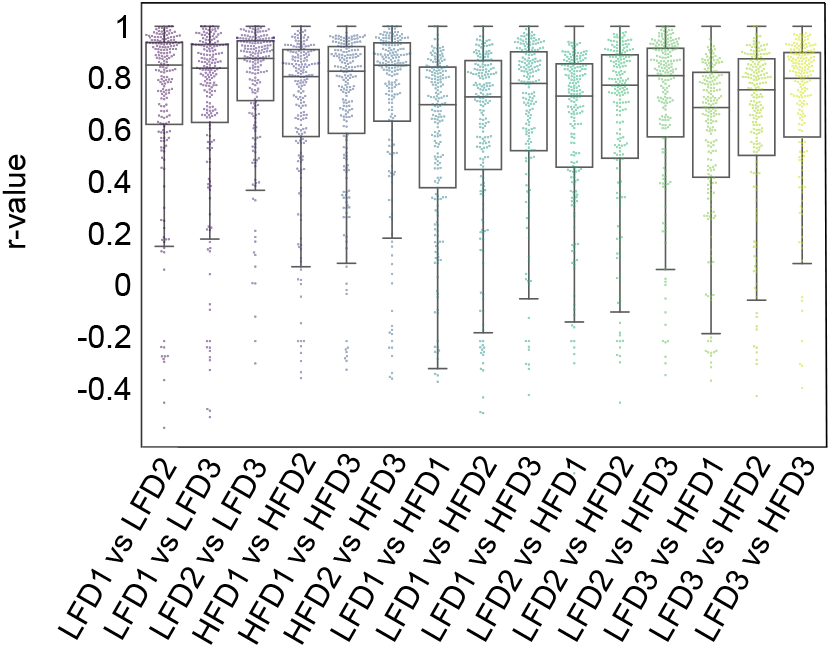
Murine liver PCP profiles indicate high reproducibility of sample replicates. Boxplot indicating correlation of PCP profiles within the indicated biological replicates of the same or different conditions.

**Figure S2:**
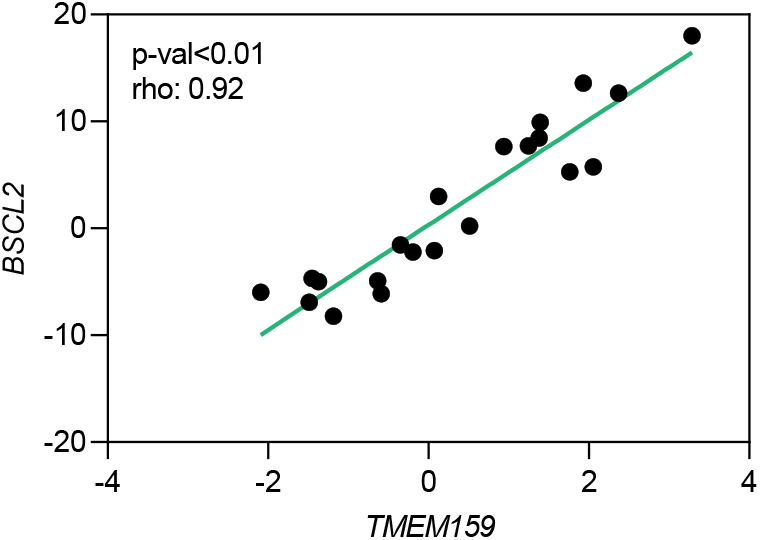
*BSCL2* and *TMEM159* knockdowns result in similar LD morphology. The LD phenotypes of *BSCL2*- and *TMEM159*-depleted cells were compared using simple regression analysis across the filtered image parameters.

**Figure S3:**
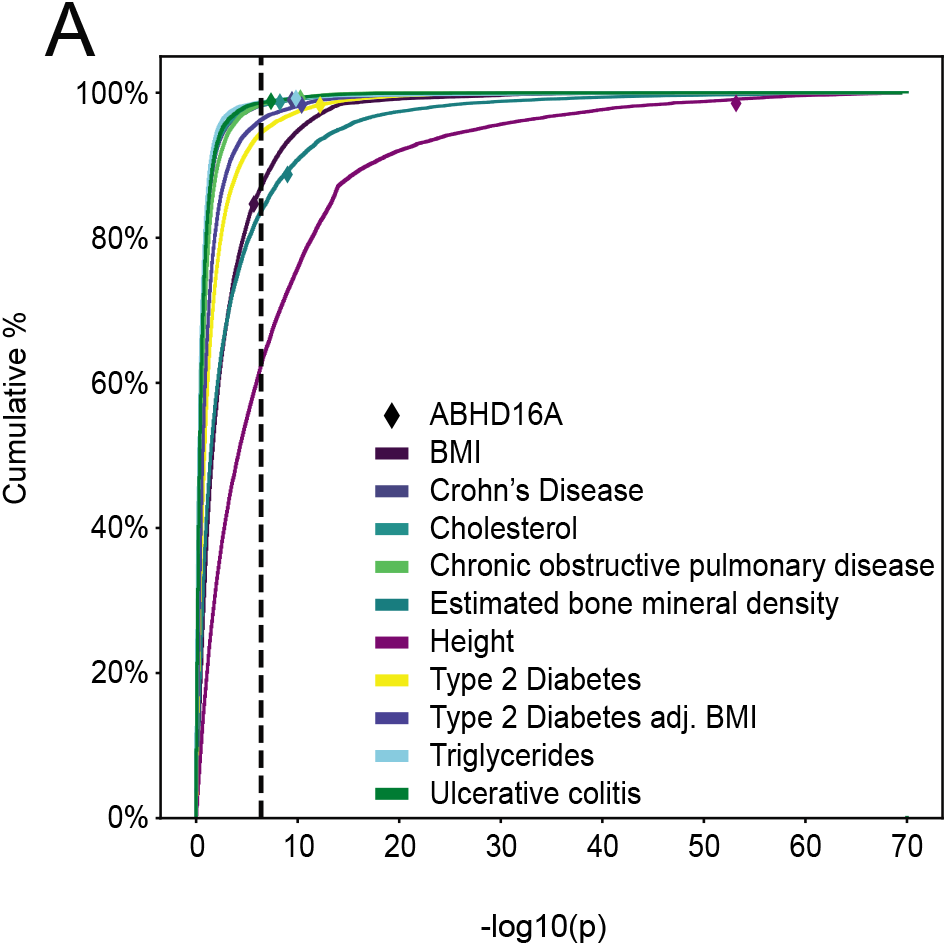
ABHD16A is significantly associated with 8 human traits. Cumulative histogram of the −log10 p values for all genes for 10 selected traits. Significance cutoff (p = 2.5e-6) is represented by the black dotted line. ABHD16A trait percentile is marked with a diamond. Abbreviations: ALT, alanine transaminase; BMI, body mass index; CAD, coronary artery disease; CHOL, cholesterol; ChildObesity, child obesity; FA160, palmitic acid; FA180, Palmitoleic acid; FA181n9, oleic acid; FG, fasting glucose; FI, fasting insulin; HDL, HDL cholesterol; IBD, Inflammatory bowel disease; LDL, LDL cholesterol; T2D, type 2 diabetes; TG, triglyceride; VAT, visceral adipose tissue volume; WAIST, waist circumference

**Table S1.** LD proteome integration

**Table S2.** Pair-wise correlation table of RNAi screen hits

## STAR METHODS

### RESOURCE AVAILABILITY

#### Lead contact

Further information and requests for resources and reagents should be directed to and will be fulfilled by the lead contacts, Robert V. Farese Jr. and Tobias C. Walther.

#### Materials availability

This study did not generate new unique reagents.

#### Data and code availability

The datasets integrated in the LD-Portal are available at https://www.lipiddroplet.org/.

### EXPERIMENTAL MODEL AND SUBJECT DETAILS

#### Cell studies

THP-1 monocyte/macrophage and SUM159 cell-culture conditions are described in Mejhert *et al*. [6]. For induction of lipid storage, cells were incubated in the presence of ac-Lipo (A6961, PanReac Applichem), ac-LDL (BT-906, Alfa Aesar), or OA (O1383, Sigma-Aldrich). Human ac-Lipo was acetylated as described [33] and OA was complexed with essentially fatty acid–free BSA (A6003, Sigma-Aldrich) at a fatty acid/albumin molar ratio of 3:1.

#### Animal studies

Mice were handled as described in Krahmer *et al*. [10]. In brief, 4-week-old C57BL/6J mice were fed either a low-fat (D12331, Research Diets) or high-fat diet (D12329, Research Diets) for 12 weeks. In accordance with an approved protocol (Animal Protection Institute of Upper Bavaria 55.2-1-54-2532-164-2015), mice were sacrificed in an *ad-libitum*-fed state, and the livers dissected for proteomic analyses.

### METHOD DETAILS

#### RNA isolation and sequencing

Total RNA isolation and sequencing procedures are described in Mejhert *et al*. [6]. In brief, total RNA was isolated from THP-1 macrophages using the QIAshredder and RNeasy Mini kits (79656 and 74106, QIAGEN). Samples were submitted to the Genomics Core at Tufts University for RNA sequencing. After quality controls and library preparation, samples were sequenced on a HiSeq 2500 using V4 chemistry (Illumina). Data analyses are described under “Processing of RNA sequencing data” below.

#### Lipid extraction and thin layer chromatography

Details on lipid extraction and thin layer chromatography are described in Mejhert *et al*. [6]. In brief, lipids were extracted from THP-1 macrophages incubated with ac-Lipo or OA using Folch’s extraction [34]. Lipids were subsequently separated by thin layer chromatography using a neutral lipid solvent (heptane/isopropyl ether/acetic acid, 60:40:4, v/v/v) as described [35] and detected by cerium molybdate staining. Quantifications were performed in Fiji [36]. For cholesterol ester quantifications, lipids were collected from lower organic phase and separated by TLC using a hexane:diethyl ether:acetic acid (80:20:1) solvent system. TLC plates were exposed to a phosphor-imaging cassette overnight and revealed by Typhoon FLA 7000 phosphor imager. Band intensities were quantified using Fiji.

#### Organellar proteomics

Mouse liver protein correlation profiles were generated as described in Krahmer *et al*. [10]. THP-1 and SUM159 LD proteomes were generated and described as in Mejhert *et al*. [6].

#### Genome-wide and secondary RNAi screens

The THP-1 macrophage RNAi screen was completed and described in Mejhert *et al*. [6]. In brief, the screen was run with samples in triplicate using an siRNA library comprising 18,119 target genes with 4 oligos per target gene. THP-1 cells were plated and differentiated for 1 day in RPMI medium containing phorbol 12-myristate 13-acetate, then transfected using Lipofectamine RNAiMAX. Subsequently, cells were grown in serum-free RPMI medium for 3 days, followed by incubations with 25 μg/mL of ac-Lipo for 2 days except for controls not containing lipids. To stain LDs and nuclei, cells were fixed with 4% paraformaldehyde and then incubated with Hoechst and BODIPY stains. 7 images per well were acquired for each channel using an Opera High Content microscope (PerkinElmer). To extend the genome-wide RNAi screen, a secondary screen was performed. Genes were randomly selected and re-screened using pools of 4 siRNAs. Cells were incubated with OA or ac-LDL to induce storage of TGs or CEs, respectively. Results were generated as described above, and results were compared among the genome-wide RNAi screen and the validation studies by performing pair-wise correlations using the set of selected image features. Data analyses are described under “Image analyses” below.

#### Structure predictions

To predict transmembrane helices potentially present in C16orf54, the TMHMM Server v. 2.0 was used with default settings [37]. The FASTA sequence of C16orf54 was obtained from UniProt database [38].

#### Cholesterol esterification assays

Cholesterol ester formation was measured in cells and lysates. For both assays, samples from RISC-free control or siMSRB3 transfected macrophages were included, and pharmacological inhibition of cholesterol ester formation was used as an assay control (S9318, Sigma-Aldrich). The assay performed in cells was originally described in [39]. In brief, cells were starved from serum for 3 days. 16 hours prior to performing the assay, compactin and mevalonate were added to the cell-culture media to inhibit endogenous cholesterol biosynthesis. Cells were subsequently incubated with the indicated amount of ac-LDL for 7 hours. To determine cholesterol esterification levels, [^14^C]oleate was added to the media for the last 2 hours. After thorough washes with cold PBS, lipids were extracted and quantified as described under “Lipid extraction and thin layer chromatography”. The *in vitro* acyl CoA:cholesterol acyltransferase (ACAT) activity in cell lysates was measured as described [40] with some modifications. In brief, THP-1 cells were lysed in lysis buffer (50 mM Tris Cl, pH 7.4, 250 mM sucrose, with protease inhibitors (11873580001, Roche)). Cell pellets were resuspended in ice-cold lysis buffer and sonicated using ultrasonic homogenizer (Biologics, Inc., model 3000MP) for 10 sec with 30% amplitude. Cell homogenate was centrifuged at 3000 x g at 4°C for 5 min and supernatant was used as enzyme source. Total ACAT was measured at Vmax substrate concentrations. Assay mixture contained 20 μg of proteins, 200 μM of cholesterol (dissolved in ethanol), 25 μM of oleoyl-CoA, which contained [^14^C] oleoyl-CoA as tracer, and 1 mM MgCl_2_ in an assay in buffer containing 100 mM Tris-HCl (pH 7.4) and protease inhibitors. Total reaction volume was 200 μl, and the reaction was performed in 2 ml tubes. The reaction was carried out for 30 min at 37°C in a water bath with shaking. Reaction was stopped by adding 1 ml chloroform and methanol (2:1) and acidified water (2% orthophosphoric acid). After stopping the reaction, tubes were vortexed well and centrifuged 10,000 x g at room temperature for 10 min. Quantifications were performed as described under “Lipid extraction and thin layer chromatography”.

#### SDS page and western blot

Details on SDS page and western blotting, are described in Mejhert *et al*. [6]. Primary antibodies targeting calnexin (sc-46669, Santa Cruz Biotechn.) and ACAT1 were used. The latter was kindly provided by Drs. Ta-Yuan and Catherine Chung-Yao Chang (Department of Biochemistry, Dartmouth Medical School).

### QUANTIFICATION AND STATISTICAL ANALYSIS

#### Statistics

Visualizations and statistical analyses of results were performed using appropriate packages in RStudio (version 1.0.143) as described under each subheading.

#### Processing of RNA sequencing data

Raw sequencing data were analyzed as described in Mejhert *et al*. [6]. Briefly, transcript abundance was quantified using Salmon [41], results were imported into RStudio using tximport [42], and differentially expressed genes were identified using DESeq2 [43]. Gene set-enrichment analysis was performed to identify gene sets regulated by lipid storage. For this, the Hallmark track was used from the Molecular Signatures Database. As a proxy for SREBP2 and NFκB activity, pathways were created based on target genes identified in published studies [44, 45]

#### Processing of proteomic data

Details on the processing of proteomic data are described in Mejhert *et al*. [6] and Krahmer *et al*. [10]. In brief, correlation profiling was applied to map the cellular localization of proteins and phosphopeptides in mouse liver. Cellular localizations are assigned by support-vector machine-based learning on the generated profiles. For THP-1 and SUM159 data, fold changes comparing LD fractions with total cell lysates were based on label-free quantification. To calculate 99% confidence intervals for canonical LD proteins, the top 50 high-confidence proteins targeting to LDs were extracted [12] and overlapped with the results presented in this study.

#### Image analyses

Details on the image analyses were described in Mejhert *et al*. [6]. In brief, CellProfiler was used to extract features from the images. For each extracted image feature, the median rz-score was calculated per gene. Image feature replicates were compared pairwise across the screen, and non-reproducible parameters were excluded. After this, a correlation matrix was generated by correlating all included image features with each other, and the dimensionality of the matrix was tested using hierarchical clustering. Features were excluded if they covaried, and genes were classified as hits if they were distributed top/bottom 15 for one image parameter and/or top/bottom 50 for more than one of the remaining high-confidence image parameters. The RNAi screen hits were pairwise correlated, based on the filtered image features, and the resulting matrix containing Spearman’s rho values was clustered using the pheatmap package (with default clustering methods and cutree_rows/cols set to five). All steps downstream of the CellProfiler analysis were performed in RStudio. The top-three and top-10 neighbors for proteasome subunits and *BSCL2* were extracted and highlighted using Cytoscape or the pheatmap package, respectively. For images from THP-1 macrophages transfected with control or *C16orf54*-targeting siRNAs, the filtered features from the RNAi screen were extracted, z-scored and clustered using the pheatmap package.

#### Expression analyses of FANTOM CAT browser data

*C16orf54* expression levels across 571 human cells and tissues were extracted from the FANTOM CAT browser [46]. Samples were ranked from high to low abundance of *C16orf54* expression levels.

#### Human genetic association analysis

Genetic association results in the LD-Portal are derived from the Common Metabolic Diseases Knowledge Portal (CMDKP; cmdkp.org), a public resource that aggregates genetic association results for >300 metabolic diseases and traits. In the CMDKP, genetic association results are meta-analyzed using the METAL algorithm, accounting for sample overlap between datasets, to generate “bottom-line” single-variant associations for each disease and trait. These are then analyzed with the MAGMA method (using default parameters) to generate gene-level association scores for each trait and each gene [28]. MAGMA gene-level association, scores are calculated based on the average association Z-scores for SNPs within a fixed window of the gene, after correcting for correlations among single-nucleotide polymorphism.

### ADDITIONAL RESOURCES

Lipid droplet knowledge portal: https://www.lipiddroplet.org.

